# In-Bead spheroid formation of dental pulp stem cells in alginate microbeads co-encapsulating NecroX-5-loaded nanoparticles for enhanced cell survival

**DOI:** 10.64898/2026.06.14.732109

**Authors:** Floriane Debuisson, Bernard Ucakar, Kevin Vanvarenberg, Catherine Le Visage, François Loll, Ana Santos-Coquillat, Ariane Mwema, Anne des Rieux

**Affiliations:** UCLouvain, Louvain Drug Research Institute, Advanced Drug Delivery and Biomaterials, 1200, Brussels, Belgium; Nantes Université, ONIRIS, INSERM, Regenerative Medicine and Skeleton, RMeS, UMR 1229, F-44000 Nantes, France

**Keywords:** mesenchymal stem/stromal cell, microfluidics, alginate, spheroids, NecroX-5, micro beads, dental stem cells, nanomedicines, cell viability

## Abstract

Mesenchymal stem/stromal cells (MSC) are key players in regenerative medicine due to their immunomodulatory properties and ability to promote tissue repair. However, their therapeutic efficacy is often limited by rapid clearance following transplantation. MSC spheroids have shown enhanced functional properties, and we hypothesize that encapsulating them within hydrogel microbeads could offer additional protection and improve their viability. We developed a continuous, single-emulsion droplet-based microfluidic protocol that forms MSC spheroids in situ within alginate droplets overnight (In-Bead), before cross-linking and bead retrieval. Unlike double-emulsion approaches, the single-emulsion format yields homogeneously distributed nanomedicines, and the short, cell-permissive cross-linking produces ultra-soft, tissue-mimetic (∼150 Pa) beads that remain retrievable. SCAP were successfully encapsulated within 275 µm alginate microbeads, forming spheroids of approximately 80 µm in diameter. Encapsulated SCAP spheroids retained their immunomodulatory properties. The process was further optimized by incorporating nanomedicines into the alginate solution before the formation of droplets, thus forming hybrid beads (Sph.Beads/NP). Nanomedicines were loaded with NecroX-5, a necrosis inhibitor, to improve SCAP viability further. Live/Dead assays indicated a protective effect of the nanomedicines, supporting the potential of this system for advanced cell delivery in regenerative applications.

**Graphical abstract:** A combination strategy enhancing MSC viability through spheroid formation, microencapsulation, and nanomedicine association achieved by microfluidic encapsulation with In-Bead spheroid formation. *Created with BioRender*

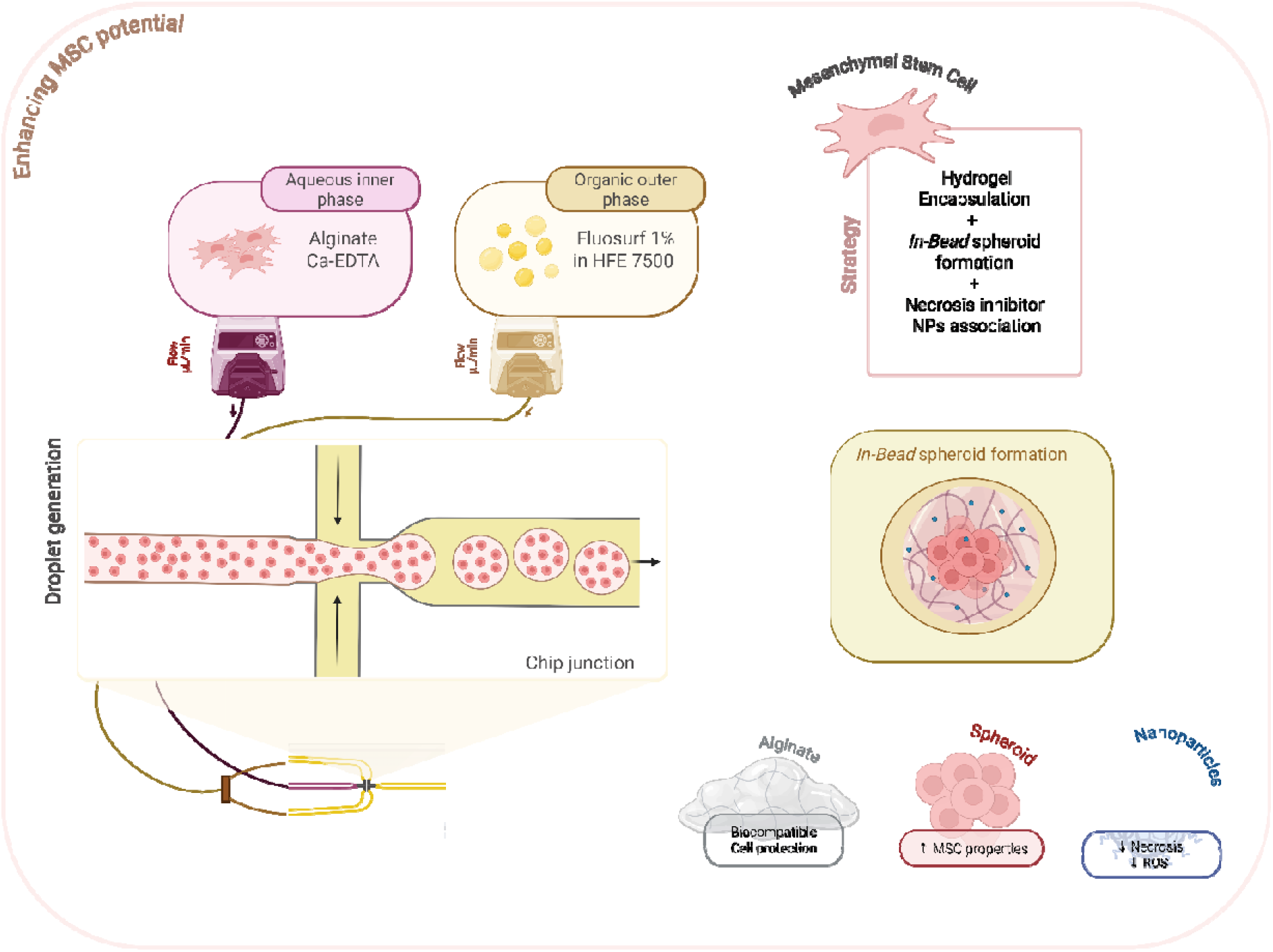

## 1 INTRODUCTION

Mesenchymal stem/stromal cells (MSC) play a central role in regenerative strategies, both in tissue engineering and cell therapy, due to their ability to support tissue regeneration [2] and modulate the local microenvironment through their secretome [3]. Among alternatives to bone marrow-derived MSC, dental stem cells, particularly stem cells from the human apical papilla (SCAP), have garnered growing attention [4]. SCAP are especially promising for neuroregeneration, as they originate from the neural crest [5] and have been studied for their potential in spinal cord injury repair [6, 7].

However, the therapeutic efficacy of MSC remains limited, primarily due to poor survival and low retention following transplantation. Different strategies have been developed to improve MSC survival after transplantation [11]. While MSC spheroid formation, hydrogel encapsulation, and nanomedicine-based exogenous support have each been explored as independent strategies to enhance cell survival and function, their integration represents a novel and particularly promising avenue. By combining these approaches, it may be possible to leverage the enhanced immunomodulation and stemness of spheroids [12–14], the controlled microenvironment and immune protection of hydrogels [15, 16], and the targeted, sustained delivery enabled by nanomedicines [17].

Several biomaterials have been used to protect MSC. Among these, alginate is one of the most widely studied materials [26]. Alginate cross-links via interactions with divalent ions, primarily calcium (Ca^2+^), with gelation triggered by CaCl_2_ addition, pH-dependent CaCO_3_ dissolution, or chelation with EDTA [27, 28]. Further enhancement of MSC properties and post-transplantation survival can be achieved through exogenous support, such as the co-administration of bioactive molecules. This includes growth factors typically delivered via hydrogel co-encapsulation or micro/nanocarrier association to protect them and enable their sustained release. Among the critical challenges to MSC survival, necrosis plays a major role.

NecroX-5, a potent necrosis inhibitor, mitigates oxidative cell death by scavenging reactive oxygen species (ROS) [34, 35]. Building on this, we previously optimized encapsulation of NecroX-5 in polymeric nanoparticles (NP), which provided prolonged necrosis inhibition, thereby improving cell viability [17, 36]. We therefore hypothesized that combining MSC with NecroX-5-loaded nanomedicines, particularly within the protective microenvironment of alginate hydrogels, could enhance cell survival by addressing both oxidative damage and the hostile post-transplantation environment.

While we previously proposed the combination of MSC spheroids with NP using a magnetic bioprinting technique [17], the association efficiency between cells and NP remained suboptimal. We then explored another strategy aiming at co-encapsulating MSC spheroids and NecroX-5 NP in alginate microbeads by microfluidics. Droplet-based microfluidics is a technology that precisely manipulates fluids at the microscale and can be used to generate hydrogel beads [37]. Unlike conventional methods such as electrostatic extrusion, microfluidics offers unparalleled control over droplet formation, yielding hydrogel microbeads with uniform size, high throughput production, and tunable cross-linking dynamics. This level of precision ensures a consistent and reproducible microenvironment, critical for optimizing cell viability and function [38]. While droplet-based microfluidics is widely established for single-cell encapsulation, its application to spheroid-NP co-encapsulation remains underexplored, highlighting the novelty of our approach [39].

Unlike conventional approaches, where spheroids are pre-formed before encapsulation, our method enables the formation of exactly one spheroid in pre-gel alginate droplets (In-Bead), followed by controlled cross-linking. This ensures optimal bead structural integrity while preserving spheroid viability, addressing a key limitation of existing techniques. Additionally, we investigated the feasibility of co-encapsulating NecroX-5-loaded NP within these alginate beads and evaluated their impact on MSC survival and function.

## 2 MATERIALS & METHODS

### 2.1 Materials

Low viscosity alginic acid sodium salt extracted from seaweed was purchased from Sigma Aldrich (no. 180947, lot MKCS7881 with M/G ratio 1.56, molecular weight 12-80 kDa, 17 cP for 1% in H_2_O). NecroX-5 (methanesulfate) and NecroX-2 were purchased from Enzo Life Sciences (New-York, NY, USA). Fluosurf 2% and HFE 7500 were purchased from Unchained Labs (Pleasanton, CA, USA). PFO (1H,1H,2H,2H-Perfluoro-1-octanol) was purchased from TCI Chemicals (Tokyo, Japan). Calcein-AM was purchased from Biotium (Fremont, CA, USA). Resomer 502H® (D, L-lactide-co-glycolide) (PLGA) was a kind gift of EVONIK (Essen, Germany). DiD’ solid; DiIC18(5) solid (1,1’-Dioctadecyl-3,3,3’,3’-Tetramethylindodicarbocyanine, 4-Chlorobenzenesulfonate Salt), TRIzol® Reagent, FluoReporter™ Blue Fluorometric dsDNA Quantitation Kit, and Ethidium homodimer D-1 were purchased from inVitrogen, Thermo Fischer Scientific (Waltham, MA, USA). Penicillin/Streptomycin (PEST), HEPES (1M) buffer, and StemPro™ Accutase™ were purchased from Gibco™, Thermo Fischer Scientific (Waltham, MA, USA). GoScript Reverse transcriptase and GoTaq® qPCR Master Mix were purchased from Promega (Madison, WI, USA). Poly(ethylene glycol) methyl ether-block-poly(lactide-co-glycolide) (PEG-PLGA – PEG average Mn 5,000, PLGA Mn 7,000, ratio 0.7), Sodium Cholate hydrate, EDTA calcium, Minimum Essential Medium (MEM) Eagle alpha, Fetal Bovine Serum (FBS), Ammonium formate LiChropur™ LC/MS, HiPerSolv CHROMANORM Formic acid 99% LC/MS were purchased from Sigma Aldrich (St. Louis, MO, USA). Recombinant Human TNF-α, Recombinant murine TNF-α, Recombinant Human IFN-γ, and Recombinant murine IFN-γ were purchased from PEPROTECH (London, UK).

### 2.2 Formulation of SCAP spheroids in alginate microbeads

#### 2.2.1 Cell-loaded alginate droplet generation by flow-focusing microfluidics

Dental stem cells from human apical papilla (SCAP) (RP89 cells [40]) were obtained from Prof. Diogenes (San Antonio, Texas). SCAP were registered in UCLouvain Biobank (BB190051). SCAP were cultured in alpha-MEM medium supplemented with 10% FBS and 1% PEST. SCAP expressing Green Fluorescent Protein (GFP+) and luciferase (Luc+) were obtained by lentivirus transduction, kindly performed in the Michiels’ lab (de Duve Inst., UCLouvain), as previously described [17, 41, 42].

For bead production, SCAP were retrieved (Accutase™) and resuspended at 5.10^6^ cells/mL in alginate (0.75% w/v, charcoal-treated) and Ca-EDTA (37.5 mM) in complete cell culture medium. Using a Dolomite™ microfluidics system (Unchained Lab) connected to a fluorinated Droplet Junction Chip (190 µm; P/N: 3200506), alginate droplets containing isolated cells were produced using a continuous organic phase of HFE 7500 supplemented with Fluosurf^®^ 1%. Flow rates were set at 20 µL/min for the oil phase and 10 µL/min for the alginate/cell phase. Magnetic stirring was applied to the alginate/cell mix (100 rpm) to maintain the SCAP in suspension during production. Droplet generation and cell encapsulation were monitored using a high-speed microscope (Supplementary Video 1). Droplets were collected in 1.5 ml Eppendorf tubes over 5 minutes.

#### 2.2.2 Spheroid formation and alginate cross-linking

Alginate droplets containing individual cells were collected and incubated at 37°C for 18 h to allow the formation of spheroids (In-Bead formation) [43]. Then, alginate microbeads containing SCAP spheroids (Sph.Beads) were cross-linked by the addition of acetic acid (0.2 %). Beads were washed and stabilised using PFO (>98%) and cell medium containing 25 mM HEPES, 1% (w/v) Pluronic F127, and 40 mM CaCl_2_ [28]. An anti-static gun was used to facilitate phase separation. Sph.Beads were then incubated at 37°C in fresh cell medium (10% FBS) until further use. To retrieve non-encapsulated spheroids, the alginate cross-linking step was omitted. The alginate droplets containing SCAP spheroids were disrupted, and the spheroids were washed in cell medium to retrieve the spheroids.

### 2.3 Sph.Bead characterization

#### 2.3.1 Bead morphology and cell content

Alginate beads were imaged using an optical microscope EVOS XL Core (Life Technologies) before and after cross-linking. Bead and spheroid sizes were measured using ImageJ (N=5, n=10). Non-encapsulated spheroids were used to assess the spheroid yield and the number of cells per spheroid. The spheroid yield, number of spheroids produced per sample, was determined post-production after counting (optical microscope acquisition of the sample in a 24-well and merging of 40 images) (N=2, n=3). The number of cells per spheroid was quantified using the FluoReporter® Blue Fluorometric dsDNA Quantification kit according to the manufacturer’s protocol. Spheroid and SCAP in suspension were used as controls. Fluorescence was measured (Ex/Em 360-460) using a SpectraMax ID5 (Molecular Devices) (N=3, n=3).

#### 2.3.2 Cell viability

Cell viability of Sph.Beads was assessed using a Live/Dead assay. After 1, 4, and 7 days in culture, a fraction of the beads was transferred to a new plate using a sterile plastic pipette (∼200 µL, 1/5 of the sample, ∼500 beads), and the cells were stained with Calcein-AM (1 µM) and Ethidium Homodimer (4 µM) for 1 hour. Samples were analyzed by fluorescence microscopy using an LSM-980 Multiphoton confocal (ZEISS) (N=2, n=3).

#### 2.3.3 Immunomodulatory markers

Sph.Beads gene expression of immunomodulatory markers in a pro-inflammatory environment was evaluated using RT-qPCR. Sph.Beads were placed in a 24-well plate (N=2, n=3), and underwent progressive FBS depletion to achieve a final FBS concentration of 0.75%, through progressive medium changes with FBS-free medium (Pest 1%) over the course of 24 hours{Gratpain, 2024 #14028}. Sph.Beads were then treated for 48 h with human-TNFα (20 ng/mL) & IFNγ (20 ng/mL), using non-activated and activated spheroids as controls [44]. Then, Sph.Beads were placed in Lysis Matrix D tubes (MP Biomedicals, Irvine, CA, USA) containing Trizol (500 µL) and dissociated using a Precellys Evolution (60s, 2x, 5500 rpm; Bertin Technologies). Samples were then frozen at -80°C for subsequent mRNA expression analysis. Gene expression of Sph.Beads was compared to non-spheroids.

#### 2.3.4 Impact of Sph.Beads on activated microglia

The impact of Sph.Beads on the gene expression of activated BV2 cells (murine microglial cells) was assessed by RT-qPCR. BV2 cells were seeded in a 24-well plate (50,000 cells/well) and were incubated with murine TNFα and IFNγ (20ng/mL) for 16 h. Non-activated BV2 cells (identified as M0 for simplicity) and empty alginate beads were used as controls. Activated BV2 cells were incubated with Sph.Beads (equivalent of 250,000 cells) in fresh cell medium supplemented with TNFα and IFNγ for 3 days (N=3, n=4). Then, BV2 cells were washed with PBS and incubated in 200µL of Trizol before storage at -80°C for subsequent mRNA expression analysis.

#### 2.3.5 RNA extraction and real-time q-PCR

Total RNA was extracted using the Trizol/Chloroform method as described [45]. cDNA synthesis was carried out using the GoScript Reverse Transcription System. Quantitative PCR (qPCR) was performed as previously described [7], using the GoTaq SYBR Green mix on a StepOne Plus instrument and software (Applied Biosystems). Data normalisation was performed in comparison to the expression of human RPL-13 or to the expression of murine RPL-19. Primer sequences are listed in Table 1.

**Table 1.**
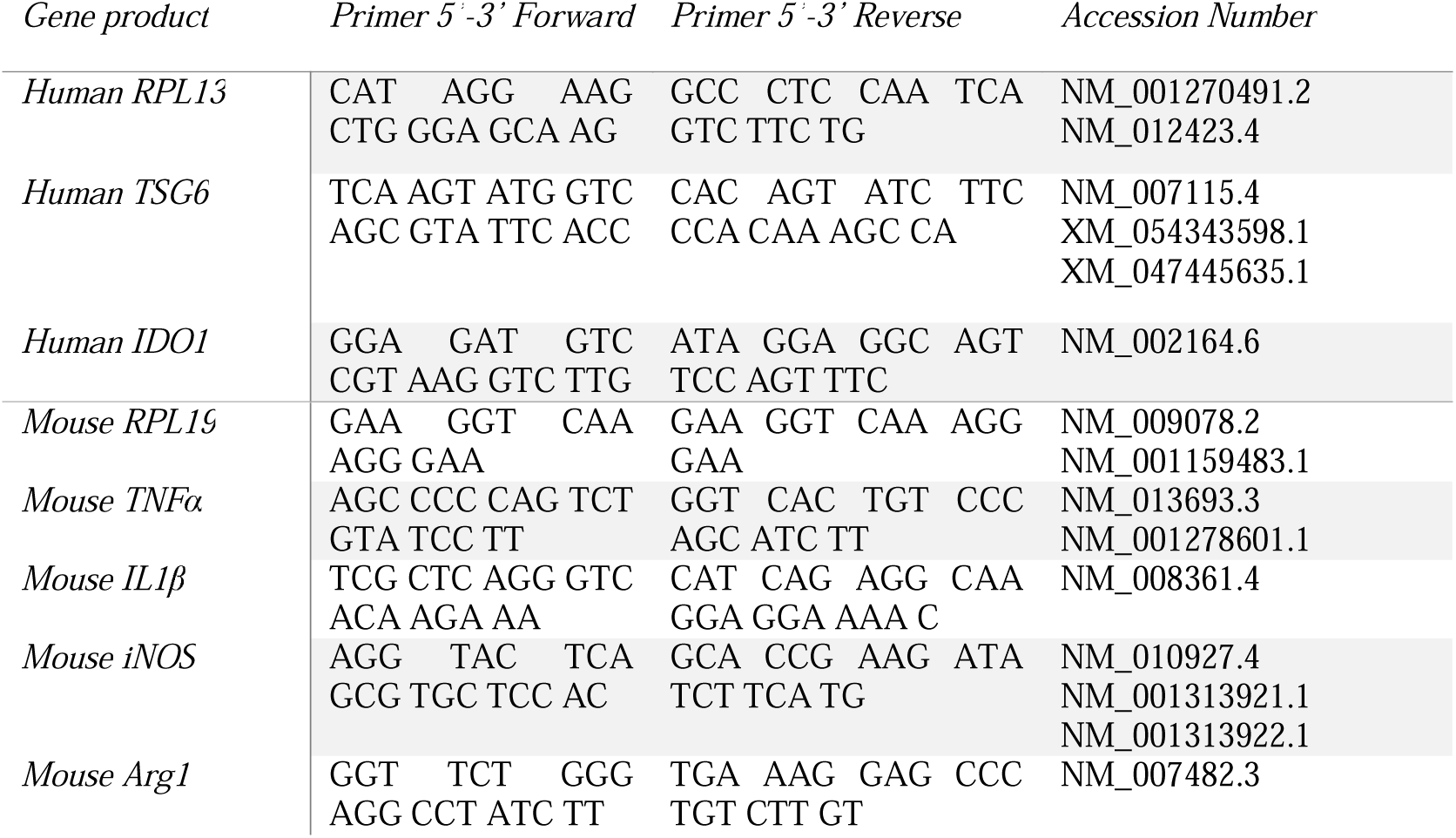
Primer sequences.

### 2.4 Formulation of Sph.Beads loaded with NecroX-5 nanoparticles

#### 2.4.1 NecroX-5 nanoparticle formulation and characterization

NecroX-5 loaded NP (Nx5.NP) were produced by single emulsion as described previously [17, 36]. Blank NP (Blk.NP) were formulated using DMSO instead of NecroX-5 solution. NP were characterized by their size, polydispersity index (PDI) and Zeta potential (ZP) by dynamic light scattering using a Zetasizer Nano ZS (Malvern Instruments). The encapsulation efficiency of Nx5.NP was measured using HPLC/UV (Shimadzu), as previously described [17].

#### 2.4.2 Production of Sph.Beads loaded with NP

Sph.Beads loaded with Nx5.NP or Blk.NP (Sph.Beads/NP; Sph.Beads/Nx5.NP or Sph.Beads/Blk.NP for Sph.Beads containing NecroX-5 NP or Blank NP, respectively) were produced as described in 2.2, with the addition of NP in the alginate/cell suspension (10µM of NecroX-5). Empty alginate microbeads (Beads), Sph.Beads and NP loaded Beads (no spheroids; Beads/Nx5.NP or Beads/Blk.NP) were used as controls.

### 2.5 Characterization of Sph.Beads/NP

#### 2.5.1 NP loading

The success of NP incorporation in the Sph.Beads was confirmed by confocal microscopy. SCAP expressing GFP+ were combined with DiD-loaded NP to form Sph.Beads/NP (Sph.Beads/DiD.NP). Live-cell nuclei staining was done using Hoechst 33342. Fluorescence microscopy images were acquired after 6, 24, and 96 hours of incubation using Z-stack acquisition (N=2, n=3).

NecroX-5 loading in the Sph.Beads/Nx5.NP was quantified by LC-MS after NecroX-5 extraction from the beads. Eighteen hours post-production, alginate droplets were collected (no cross-linking) in cell medium containing 10% FBS and 25mM HEPES. Samples were centrifuged for 10 min at 10,000g and NecroX-5 was extracted as previously described [17]. NecroX-2 was used as an internal standard (60 ng/mL) (N=3 n=2).

NecroX-5 release from the beads was studied as follows. Samples of Sph.Beads/Nx5.NP containing ∼5000 beads and ∼80 ng of NecroX-5, were placed in a Slide-A-Lyzer™ MINI Dialysis Device 2K MWCO (ThermoFischer Scientific) clipped on 2 mL tubes containing 1.5mL of cell culture medium supplemented with 10 % of FBS, 25 mM of HEPES, and incubated at 37°C (N=2, n=3). Samples were kept at -20°C until further Analysis. Then, 100 µL of sample was added to 300 µL MeOH containing NecroX-2 (40 ng/mL). Proteins were precipitated by vigorous vortexing. Then, samples were centrifuged at 14,000g for 10 min. Supernatants were concentrated by vacuum centrifugation using a CentriVap Concentrator (LabConco) (40°C – 2h). Samples were then reconstituted with 50 µL MeOH and 20 µL water, sonicated for 10 minutes, and vortexed before LC/MS-MS quantification.

LC/MS-MS quantification of NecroX-5, using NecroX-2 as internal standard, was performed by liquid chromatography coupled to tandem mass spectrometry as previously described using a LC/MS-MS 8040 (Shimadzu) set to a negative electrospray ionisation mode with a multiple reaction monitoring (MRM-) [17].

#### 2.5.2 Rheological properties

Sph.Beads/Blk.NP modulus was measured using a Microsquisher^®^ (CellScale). Negative controls were Beads, Beads/Blk.NP and Sph.Beads. To approximate physiological conditions, samples were analyzed at 37°C in PBS supplemented with CaCl_2_ (1.8mM) [46]. A 30sec compression ramp was applied to the samples using a 0.1524mm diameter microbeam and a 1mm² plate until a deformation of 20% was achieved. Young’s modulus was determined according to **Equation 1** as described by Nativel et al. [46] (n=10).

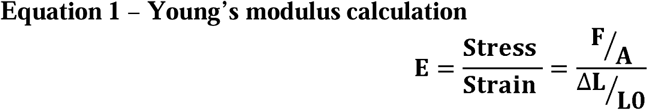

Where **E** is Young’s modulus; **F** is the force applied on a particle; **A** is the area onto which the pressure is applied; Δ**L** is the displacement, and **L0** is the initial diameter of the particle.

#### 2.5.3 Cell viability

Sph.Beads/Nx5.NP cell viability was assessed by Live/Dead assay as described in 2.3.2. Briefly, the mean green intensity, mean red intensity, and signal area of 20 spheroids were measured at different time points. Sph.Beads/Nx5.NP and Sph.Beads/Blk.NP were compared to Sph.Beads (as 100%) (N=2, n=3).

### 2.6 Statistical analysis

Statistical analyses were performed using GraphPad Prism 10 software. Outliers were identified using Grubbs’ test. The Normal distribution of the data was checked before each analysis (Shapiro-Wilk test). One-way ANOVA was used when homoscedasticity was confirmed; otherwise, non-parametric Kruskal-Wallis test was used. Statistical tests and replicates are specified in the figure legends. Statistical significance was accepted at p-value <0.05.

## 3 RESULTS

### 3.1 In-Bead formation of spheroids preserved SCAP viability and in vitro immunomodulatory properties

Our objective was to obtain one viable In-Bead SCAP spheroid per alginate microbead using microfluidics. To the best of our knowledge, this was not previously reported in the literature, so a new single-emulsion droplet protocol was established and optimized. Microfluidic alginate droplet formation was optimized to secure steady flows and reproducible droplet formation **(Supplementary video 1)**. The first challenge was to achieve homogeneous cell encapsulation throughout the production process. This was accomplished by subjecting the alginate cell suspension to a gentle stirring during the production. The bead content was monitored in real-time using the system’s high-speed camera. SCAP encapsulated in alginate droplets were incubated for 16, 18 and 24 hours to allow spheroid formation. Incubating for 18 h was the best compromise between bead stability and cell viability **(Figure 1).**

**Figure 1:**
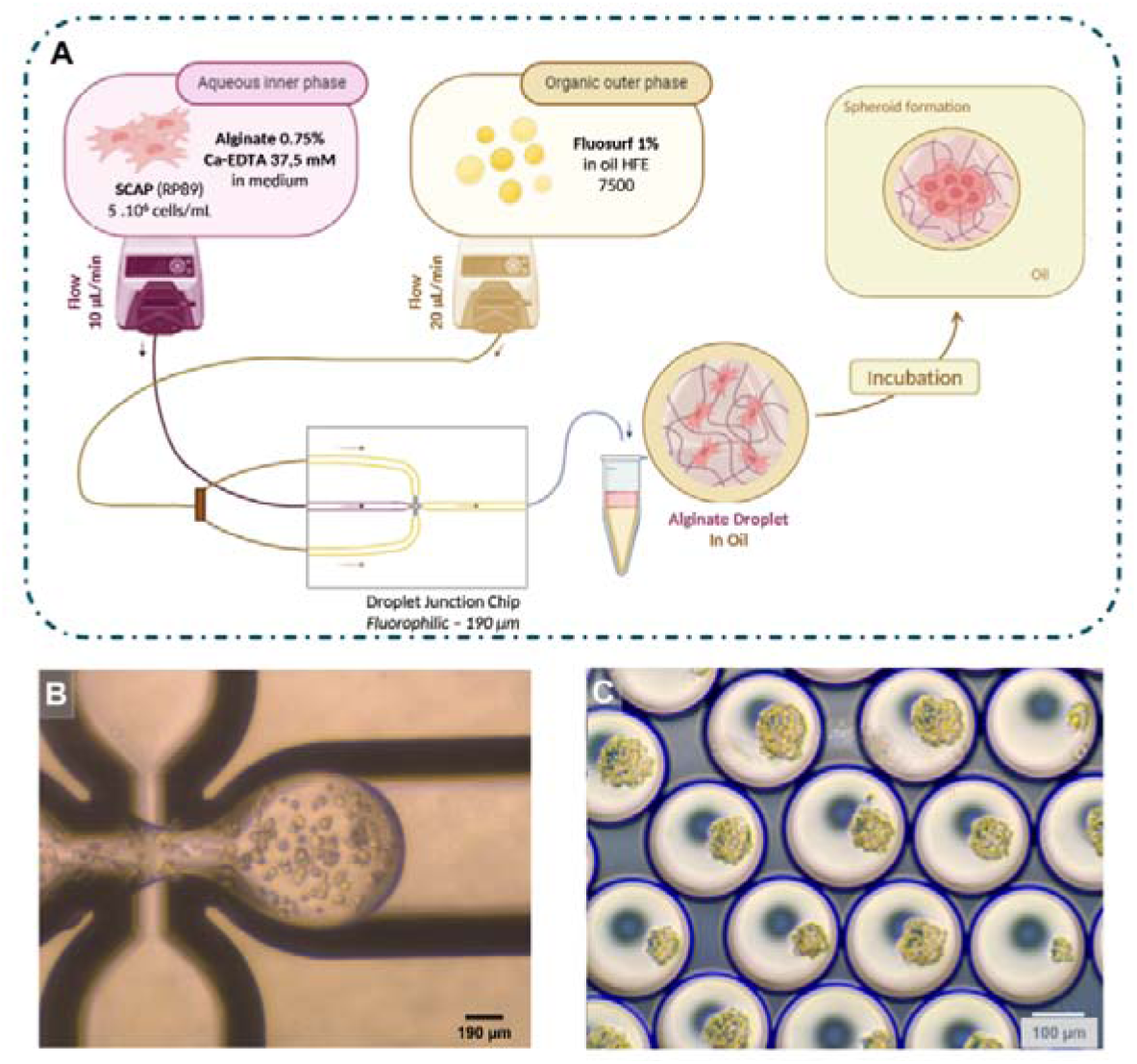
Formation of SCAP in alginate droplets using microfluidics. **A)** Optimized experimental setting. **B)** Droplet generation at the chip junction. **C)** Spheroids in alginate droplets 18 h post-incubation before cross-linking. Droplet generation was performed on a Dolomite™ microfluidic system (Dolomite Microfluidics, Cambridge, UK) using a 190 µm fluorophilic Droplet Junction Chip. Process timeline: droplet generation → 18 h incubation at 37°C → cross-linking → bead retrieval. *Created with Biorender*.

The second challenge was to achieve alginate gelation, allowing the formation of individual solid and stable beads, without compromising cell viability. Hence, alginate droplet cross-linking and bead retrieval were optimized to find a balance between exposure to a low pH and cell viability, entrapment of the spheroid, and preservation of bead morphology. Internal bead cross-linking was achieved by inducing calcium release from EDTA complexes through a pH change by the addition of acetic acid (0.2%) in the oil phase **(Figure 2A)**. HEPES was then added to buffer the solution and reduce the exposure of the cells to a cytotoxic pH. To further stabilize the alginate bead and prevent the spheroid from escaping during subsequent bead manipulations, CaCl_2_ (40 mM in 500µL) was added to the retrieval solution to induce post-acidic cross-linking. Individual Sph.Beads were retrieved by breaking the emulsion with PFO, Pluronic F127 and an antistatic gun.

**Figure 2:**
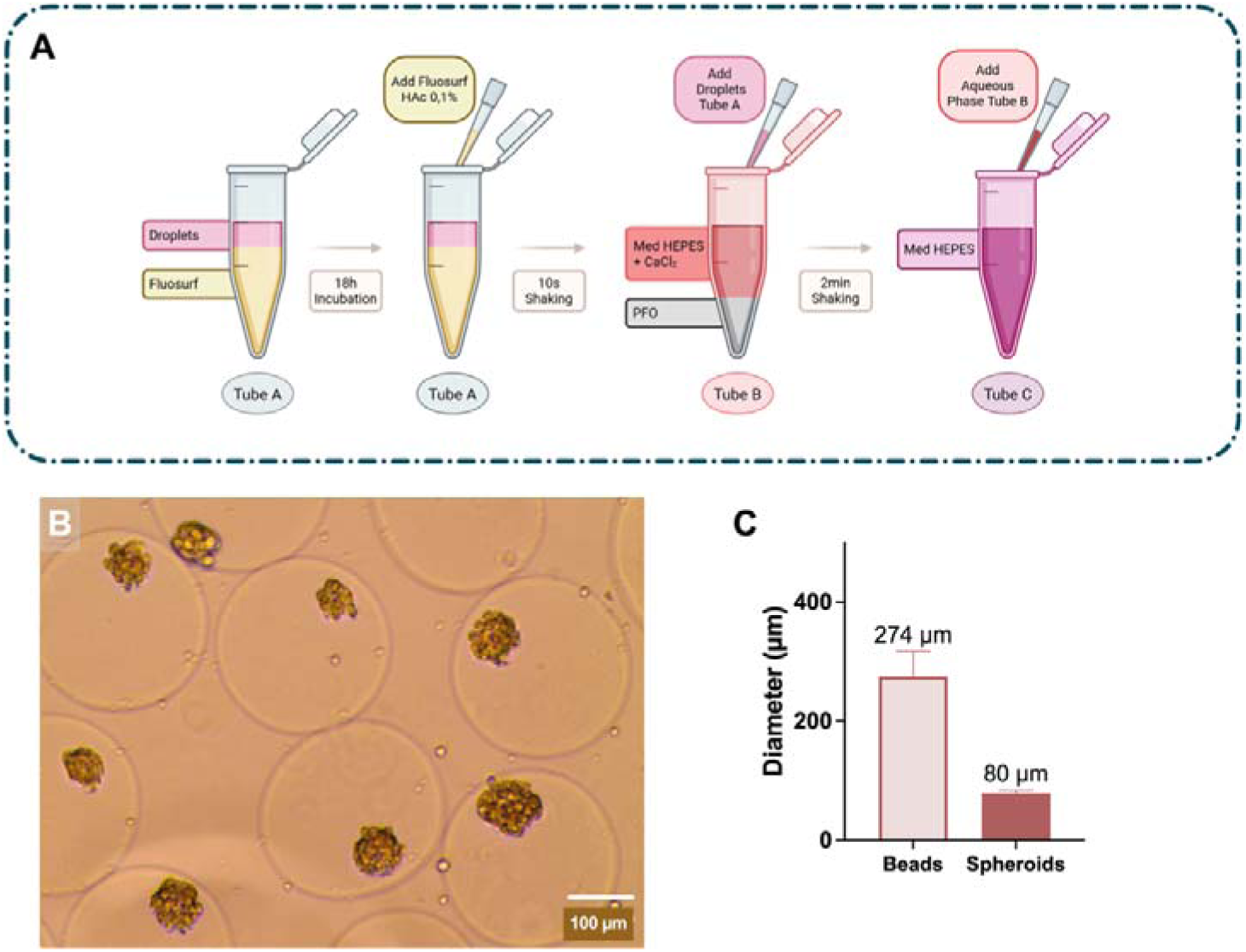
Crosslinking and retrieval of Sph.beads. **A)** O optimized experimental setting for Sph.Bead cross-linking and retrieval; Fluosurf HAc 0.2%: Fluosurf containing 0.2 % of acetic acid. **B)** Spheroids in alginate beads (Sph.Beads) after cross-linking. **C)** Bead and spheroid diameters measured with ImageJ (Mean±SD, N=5 n=10). *Created with Biorender*

This optimized protocol allowed the formation of viable Sph.Beads **(Figure 2B)**, presenting a mean diameter of 274 µm and a spheroid of a mean diameter of 80 µm composed of an average of 97 ± 24 cells **(Figure 2C)**. A theoretical number of 250,000 cells (CI 95% 225,000-300,000) was introduced into the microfluidics system for the formulation of one batch. Bead formation rate was 538 ± 79 beads/min.

Preservation of SCAP viability in Sph.Beads was confirmed using a Live/Dead assay **(Figure 3A&B)**. One day after formation, Sph.Beads showed high cell viability. Over time, viability remained high, although a slight decrease was observed after 7 days in culture. Bright-field images showed that bead integrity was preserved over time.

**Figure 3:**
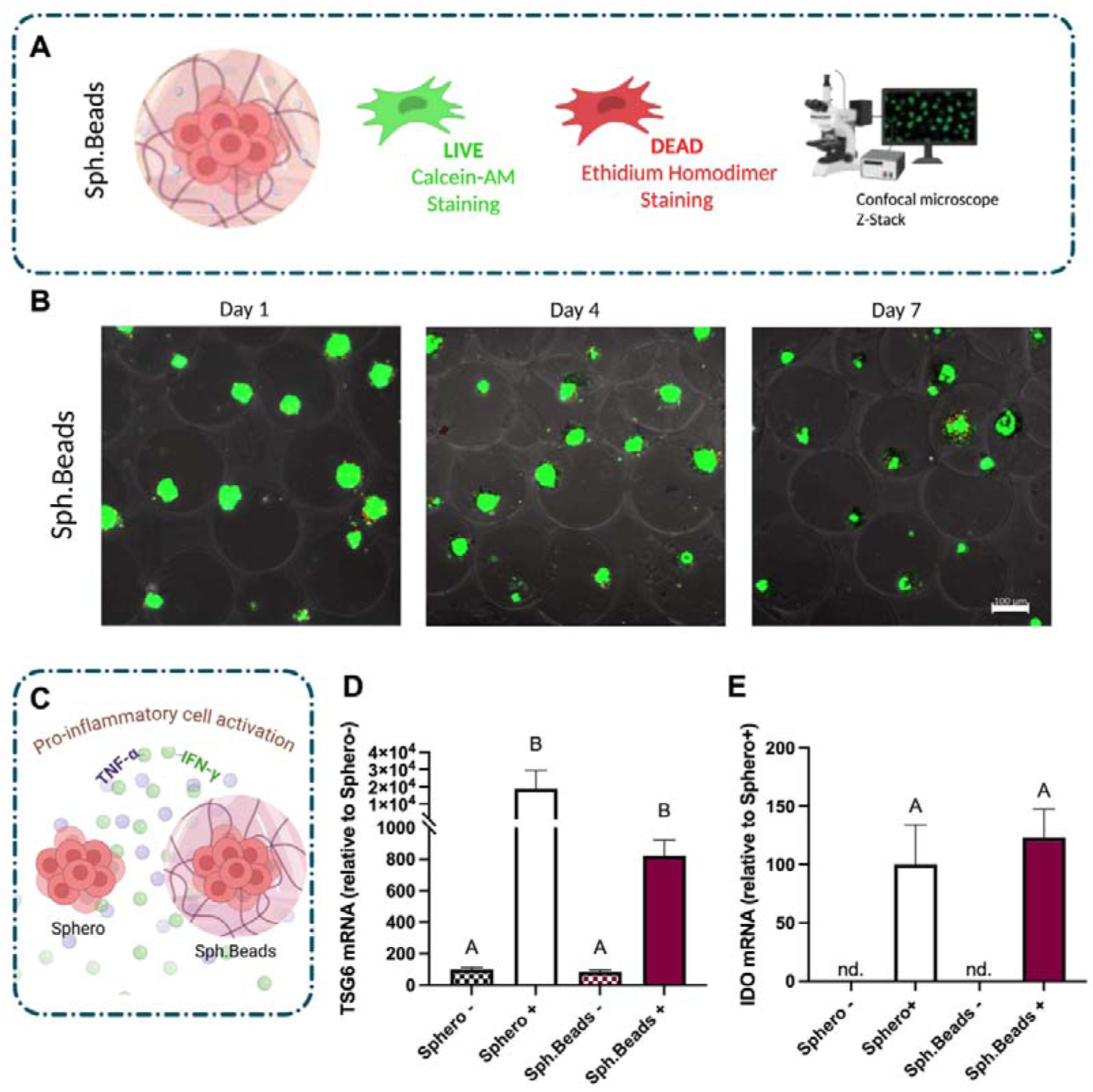
Characterization of spheroids in alginate microbeads. **A)** Experimental setting of the Sph.Beads Live/Dead assay. **B)** Sph.Beads Live/Dead assay (Scale bar 100 µm). Live cells are stained in green (Calcein AM), and dead cells are stained in red (Ethidium homodimer). Bright-field acquisition confirmed spheroid encapsulation and bead integrity over time. **C)** Experimental setting of pro-inflammatory cell activation with TNFα and IFNγ. **D)** TSG6 mRNA expression of activated (+) or non-activated (-) spheroids and Sph.Beads (Mean±SEM, N=2 n=3). Conditions not linked by the same letter are significantly different (p<0.05, Kruskal-Wallis test followed by a Dunn’s post-hoc test). **E)** IDO mRNA expression of activated (+) or non-activated (-) spheroids and Sph.Beads (Mean± SEM, N=2, n=3) (p<0.05, Mann-Whitney test). IDO and TSG6 expression were normalized to the human housekeeping gene RPL13 and expressed as fold-change relative to the unstimulated control. *Created with Biorender*

Then, the impact of a pro-inflammatory stimulus on Sph.Beads expression of immunomodulatory molecules was evaluated. TSG6 and IDO gene expression were both significantly increased upon cell activation, whether SCAP spheroids were encapsulated in alginate beads or not, indicating that encapsulation did not suppress the ability of the cells to react to a pro-inflammatory priming **(Figure 3C-E)**.

### 3.2 Sph.Beads decreased pro-inflammatory marker expression in microglial cells in vitro

The aim was to assess whether Sph.Beads maintained SCAP ability to reduce the pro-inflammatory cytokine expression of activated murine microglia cells (BV2 cells) [47, 48]. Sph.Beads were co-cultured with BV2 cells activated by TNFα and IFNγ. Activation increased the expression of pro-inflammatory cytokines (TNFα, IL1β, iNOS) in BV2 cells compared to non-activated BV2 cells (M0). TNFα expression was significantly reduced when BV2 cells were treated with Sph.Beads, but not with Beads, while IL1β expression was not impacted regardless of the treatment **(Figure 4)**. iNOS expression tended to decrease, while arginase1 (Arg1) expression, a marker of pro-resolutive microglia, tended to increase when BV2 cells were treated with Sph.Beads but it was not statistically significant.

**Figure 4:**
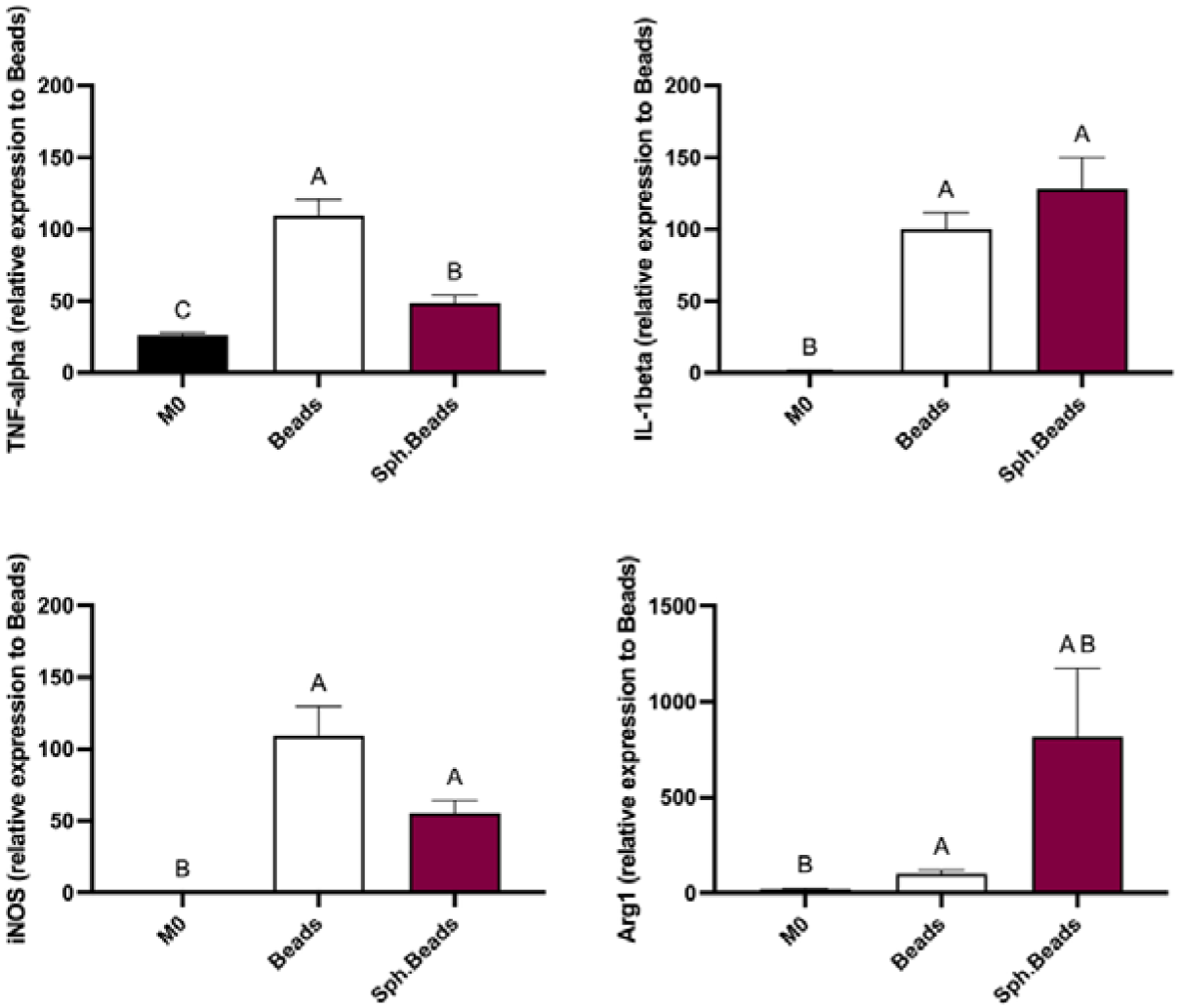
Impact of Sph.Beads on activated microglia gene expression of inflammatory markers. BV2 cells were co-cultured with Sph.Beads for 3 days in the continued presence of TNFα/IFNγ. Then, mRNA expression of pro- and anti-inflammatory markers of activated BV2 cells treated with Sph.Beads was analysed by RT-qPCR. Non-activated microglia (M0) and activated microglia treated with empty alginate beads (Beads) were used as controls. Gene expression was normalized to the mouse housekeeping gene RPL19 and expressed as fold-change relative to Beads (Mean ± SEM, N=3, n=4). Statistical differences were analysed using a Kruskal-Wallis test followed by a Dunn’s post-hoc test. Conditions not linked by the same letter are significantly different (p<0.05).

### 3.3 Addition of NecroX-5 loaded NP in Sph.Beads increased cell viability

#### 3.3.1 NecroX-5 NP physico-chemical properties

NP loaded with NecroX-5 (Nx5.NP) and blank NP (Blk.NP) exhibited similar sizes, PDI, and zeta potential (ZP) (**Table 2**). NecroX-5 was encapsulated in PLGA/PLGA-PEG NP with an encapsulation efficiency of 50%.

**Table 2.** Physicochemical characterization of NecroX-5 NP.

|  | **Nx5.NP** | **Blk.NP** |
|---|---|---|
| Size (nm) | 237 ± 26 | 238 ± 21 |
| PDI | 0.12 ± 0.04 | 0.12 ± 0.05 |
| ZP (mV) | -41 ± 2 | -40 ± 1 |
| Drug Encapsulation Efficiency (%) | 49 ± 6 | / |

Data are expressed as Mean ± SD. N ≥ 10

#### 3.3.2 Characterization of Sph.Beads/NP

The presence of fluorescent (DiD) NP in Sph.Beads was validated by confocal imaging after production (6h), showing a homogeneous distribution **(Figure 5A&B)**. The signal decreased over time, which might indicate a release of the NP or of DiD from the beads.

**Figure 5:**
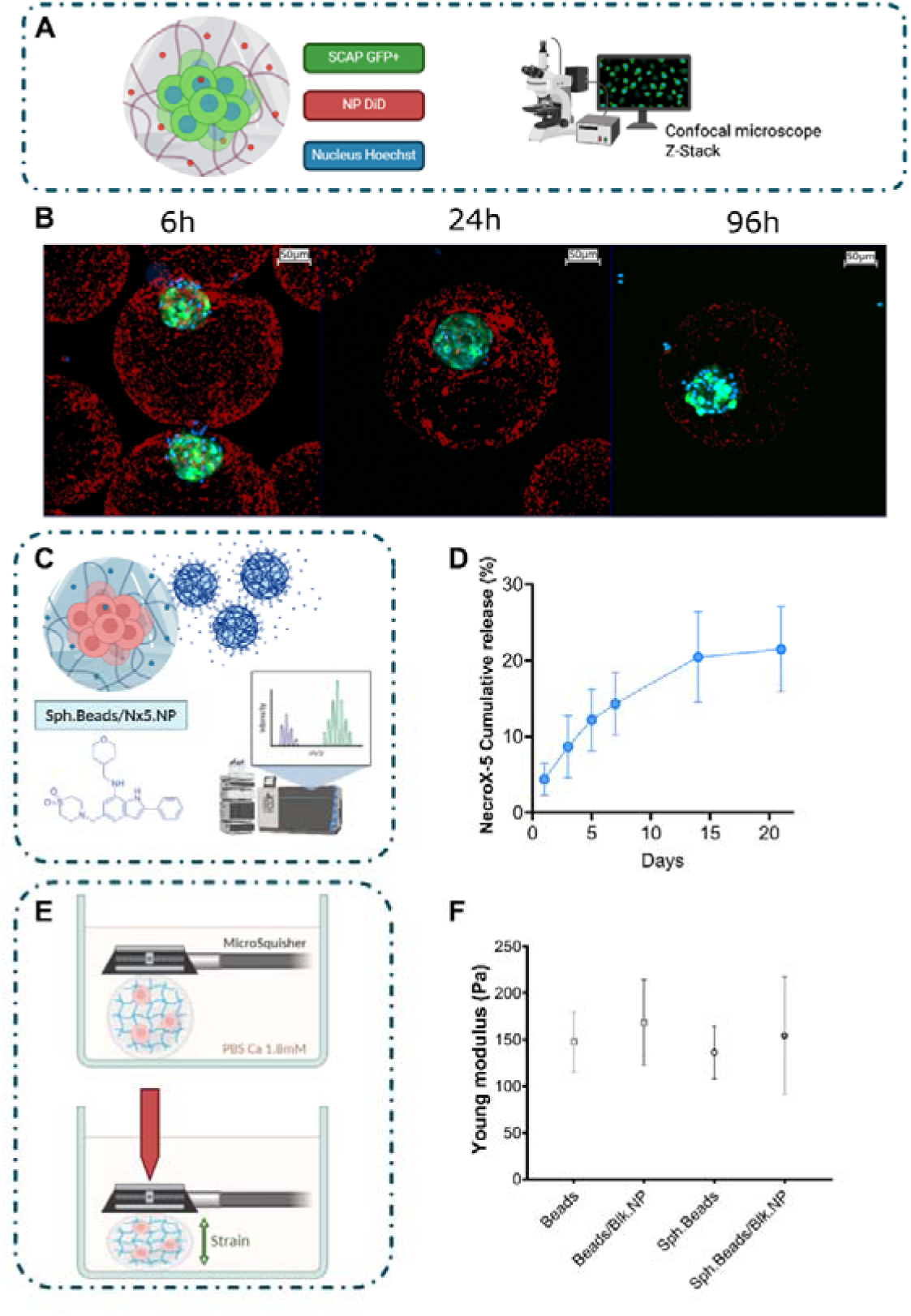
Characterization of Sph.Beads/NP. **A)** Experimental setting of confocal imaging. **B)** NP loaded with DiD (red) encapsulated in Sph.Beads formed with GFP+SCAP (green), with nuclei stained with Hoechst 33342 (blue) (Scale bar 50 µm). **C)** LC/MS-MS quantification of NecroX-5 release. Confocal imaging settings were kept constant across all time points; the decrease in fluorescence over time reflects nanoparticle release and/or dye diffusion. **D)** NecroX-5 cumulative release (in %) from Sph.Beads/Nx5.NP over 21 days (Mean±SD, N=2 n=3). **E&F)** Young’s modulus measurement (Pa) of Sph.Beads/Blk.NP using a MicroSquisher, at 37°C in PBS supplemented with 1.8 mM CaCl_2_. Beads, Beads/Blk.NP and Sph.Beads were used as controls (Mean±SD, n=10). No statistical differences were observed between the conditions (One-way ANOVA, p<0.05). *Created with Biorender*.

The loading efficiency of NecroX-5 in Sph.Beads/Nx5.NP was 24% ± 4.8%, which is equivalent to ∼2.5 µM per bead (N = 3, n =2). This concentration falls within the range reported to inhibit necrosis and modulate inflammation in vitro [34, 35, 57, 58].

A sustained release of NecroX-5 from Sph.Beads/Nx5.NP was observed, with approximately 20% cumulative release of the loaded NecroX-5 released over 3 weeks (**Figures 5C&D)**. As the initial amount of NecroX-5 in the samples was ∼80 ng, ∼16 ng were released. The results reflect the release of NecroX-5 from both the alginate beads and the NP.

The Young’s modulus of Sph.Beads/Nx5.NP was measured to assess the impact of NP and/or spheroid encapsulation on bead stiffness. The calculated modulus (∼150 Pa) showed no significant differences between conditions and thus no impact of Np or spheroids on the bead modulus **(Figure 5E&F, Supplemental Video 2)**.

#### 3.3.3 Impact of Nx5.NP on SCAP viability

The impact of Nx5.NP in Sph.Beads on SCAP viability was evaluated using a Live/Dead assay. Sph.Beads/NP, whether loaded with blank (Blk.NP) or NecroX-5 NP (Nx5.NP), showed a slight decrease in viability at Day 1 compared to Sph.Beads, but then, at Day 4 and 7, the viability of Sph.Beads/Nx5.NP was higher than the controls **(Figure 6B&C).** The ratio of living/dead cells was higher than 10, regardless of the conditions and time points **(Figure 6D)**. It increased slightly over time for Sph.Beads/Nx5.NP and Sph.Beads but not for Sph.Beads/Blk.NP. The ratio was the highest for Sph.Beads/Nx5.NP at Day 7 and was significantly higher than Sph.Beads/Blk.NP.

**Figure 6:**
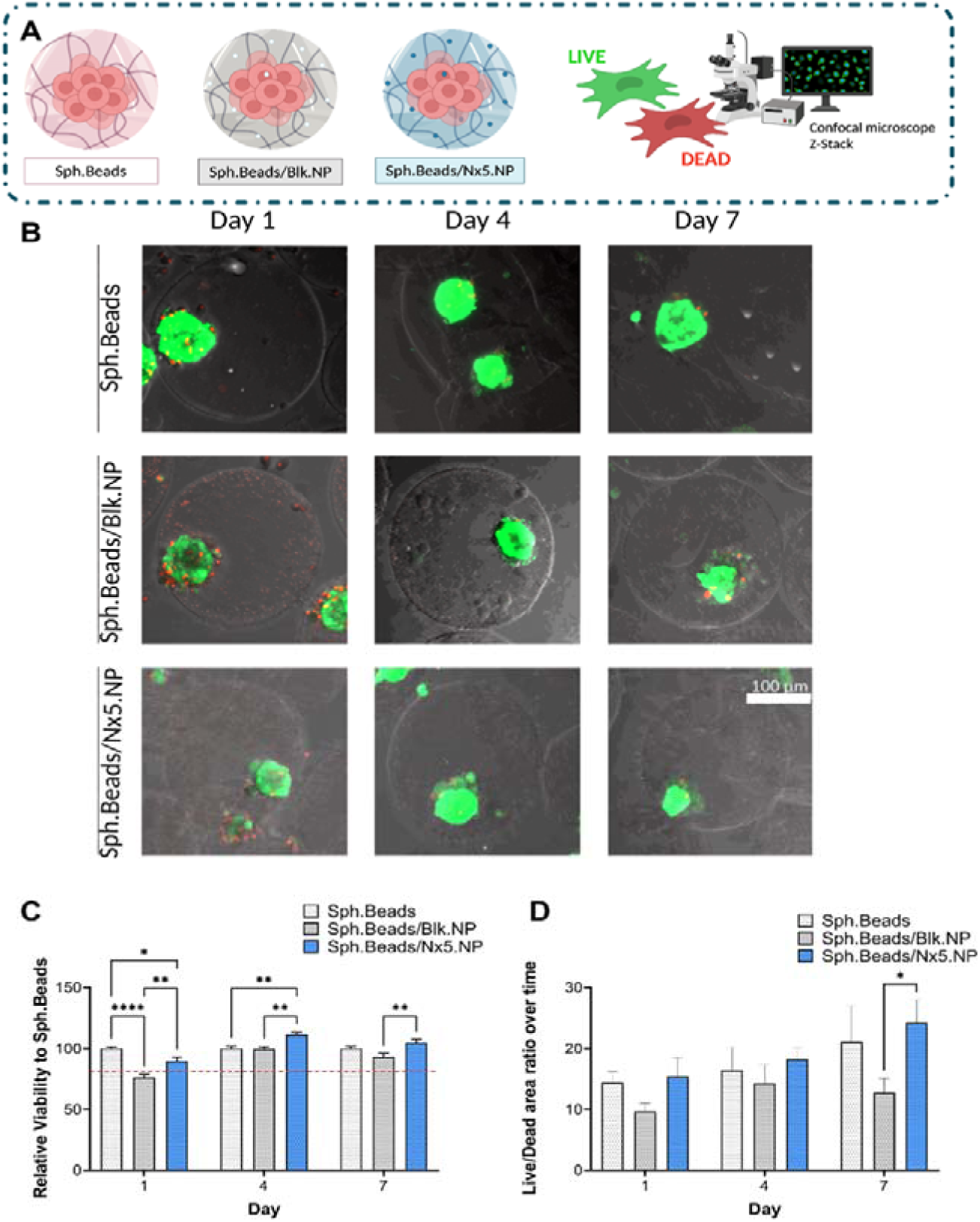
Impact of Nx5.NP on SCAP viability. **A)** Experimental setting. **B)** Live/Dead picture acquisitions by confocal imaging of Sph.Beads, Sph.Beads/Blk.NP, and Sph.Beads/Nx5.NP (Scale bar 100 µm). Confocal imaging settings were kept constant across all time points. **C)** Spheroid viability (Green signal mean intensity) normalized to Sph.Beads over time (Mean±SEM, N=2, n=3, 20 spheroids measured/condition). **D)** Ratio of living cells (area of green signal)/dead cells (area of red signal) over time (Mean± SEM, N=2, n=3, 20 spheroids measured/condition). Statistical differences were analysed using a 2-way ANOVA test followed by a Tukey’s post-hoc test. Conditions not linked by the same letter are significantly different (p<0.05). *Created with Biorender*.

## 4 DISCUSSION

Here, we developed a microfluidic hybrid delivery platform integrating in situ stem cell spheroid formation with polymeric nanomedicine incorporation, enabling the sustained local release of a protective therapeutic agent while preserving cell viability and immunomodulatory properties. We report a single-emulsion, droplet-based microfluidic platform that forms human apical-papilla MSC spheroids in situ within alginate microbeads (In-Bead), and that loads them with a necrosis-inhibiting nanomedicine (NecroX-5) in the same step to yield hybrid Sph.Beads/NP. Two design choices distinguish this platform from prior cell-in-alginate and spheroid-encapsulation work. First, the single-emulsion format, rather than the double emulsion typically required to co-load cells and drug, yields homogeneously distributed nanomedicines across the bead, avoiding the core–shell gradients of double-emulsion beads. Second, a short, cell-permissive cross-linking produces ultra-soft, tissue-mimetic (∼150 Pa) beads that remain retrievable, in contrast to the stiff beads of long cross-linking protocols. Functionally, In-Bead spheroids retained their immunomodulatory capacity, and the co-loaded nanomedicine conferred a measurable protective effect on viability in Live/Dead assays. Together, these results establish In-Bead as a scalable route to soft, drug-loaded spheroid constructs for regenerative delivery, and frame the next challenge, retaining the nanomedicine long enough to act, as the design target for the next generation of beads.

The initial step focused on optimizing the In-Bead production of spheroids (Sph.Beads) to achieve reproducible, robust formation of a single spheroid per alginate bead, while maintaining high cell viability. SCAP viability remained high throughout the production process and was sustained over one week in culture, showing that the production process was mild enough to preserve SCAP viability and that effective nutrient and waste diffusion occurred. This is consistent with the reported permeability of microbeads smaller than 500 µm [49]. This process required balancing three key factors: homogeneous encapsulation, optimal bead morphology, and spheroid retention within the bead. A delicate equilibrium had to be found between microfluidic production parameters and bead gelation and retrieval conditions to ensure both SCAP viability and retention of the spheroid within the beads. To address these challenges, we integrated several approaches to i) produce beads smaller than 300 µm in diameter, ensuring injectability in rodent models (21G needle), ii) stabilize the emulsion long enough to allow In-Bead spheroid formation (18h), iii) develop a tailored gelation protocol (combining calcium release from EDTA complexes by the addition of acetic acid, HEPES to buffer the solution and CaCl_2_ to induce post-acidic cross-linking) and retrieval to solidify beads without compromising cell viability or structural integrity (combination of PFO and medium containing Pluronic F127 and an antistatic gun). This protocol addresses all these constraints simultaneously, enabling the production of In-Bead spheroids combined with nanomedicines. While microfluidic spheroid formation has gained attraction in oncology for high-throughput screening [50, 51], the behavior of cancer cells, which can rapidly proliferate and form spheroids, differs significantly from that of MSC. This discrepancy complicates the direct translation of cancer cell protocols to MSC applications, which explains the need for protocols specifically optimized for MSC spheroid formation.

While microfluidic techniques are well established for cell encapsulation in hydrogels, including MSC [52, 53], and for spheroid formation, to the best of our knowledge, the integration of these two processes, particularly the formation of spheroids in situ alginate microbeads, remains largely undocumented. Only one group, Chan et al., described a double emulsion microfluidic protocol for MSC In-Bead spheroid formation [54]. Their method relied on custom-designed microfluidic systems and required two sequential chips to generate the double emulsion, necessitating the control of three fluid phases. These features increase operational variability and enable oil entrapment within W/O/W droplets, complicating complete oil removal compared to a single emulsion protocol. In this study, we developed a protocol using a commercial microfluidic system with a commercially available chip, providing a standardized and accessible platform for single emulsion droplet generation and cell/NP encapsulation. Our novel approach overcomes reproducibility challenges. A critical innovation was triggering external gelation without disrupting droplet integrity, enabling recovery of round, non-aggregated beads containing viable In-Bead formed spheroids.

We then demonstrated that the Sph.Bead formulation did not compromise SCAP responsiveness to a proinflammatory stimulus. Previously, we showed that SCAP, whether as single cells or spheroids (formed via magnetic bioprinting), upregulated immunomodulatory molecules such as TSG-6 and IDO upon activation with TNFα and IFNγ [17, 44]. In this study, we confirmed that the formation of In-Bead spheroids did not impair this response, as activated SCAP spheroids maintained elevated TSG-6 and IDO expression. This suggests that TNFα and IFNγ (50 and 20 kDa, respectively) effectively diffused through the alginate matrix to reach the spheroid and that encapsulated spheroids were still able to react to this stimulus [55]. Furthermore, when Sph.Beads were co-cultured with activated microglial cells, they suppressed pro-inflammatory markers (TNFα, iNOS) while upregulating the anti-inflammatory marker Arg1, indicating a shift toward an immunomodulatory phenotype.

To further enhance Sph.Bead functionality, we incorporated NecroX-5 nanoparticles (Nx5.NP) during droplet formation, producing Sph.Beads/Nx5.NP. The Nx5.NP, prepared as previously described [17], were uniformly distributed within the beads, demonstrating the reproducible loading of nanomedicines into microgels using our microfluidic protocol. A time-dependent decrease in fluorescence intensity (DiDI.NP) suggested that either NP and/or DiD was released over time from the beads. NecroX-5 release was thus confirmed, revealing sustained release kinetics. As the final Nx5.NP loading in the Sph.Beads was ∼2.5 µM, reflecting a 75% loss from the initial 10 µM concentration in the alginate stock solution, we hypothesized that during the 18-hour incubation in liquid alginate droplets enabling spheroid formation, NecroX-5 or Nx5.NP partitioned into the oily phase. Encapsulation in NP was able, however, to limit this issue, as when free NecroX-5 was encapsulated in alginate droplets under identical conditions less than 3% remained. This is consistent with findings by Guo et al., who reported a 75% burst release of Coumarin-6 from PLGA NP within 4 hours when encapsulated in PEG-alginate beads [56]. This underscores a challenge in retaining lipophilic drugs during oil-based microfluidic encapsulation. Despite this loss, the residual NecroX-5 concentration (∼2.5 µM) remained within a therapeutically relevant range, as NecroX-5 protected hepatocytes from doxorubicin toxicity at 0.125 µM [34], well below the levels typically used in vitro [57–59]. Thus, even with reduced loading, the Sph.Beads/Nx5.NP may retain biological activity.

The Young’s modulus of the Sph.Beads/NP was approximately 150 Pa, with neither In-Bead spheroid formation nor NP encapsulation significantly altering mechanical properties. Hydrogel stiffness is a critical parameter, as it influences mechanotransduction and stem cell behavior while softer gels are known to preserve stemness [60]. Our results align with findings by Cappello et al., who reported a Young’s modulus of ∼100 Pa for 200 µm beads fabricated from 1% low-viscosity alginate using microfluidics [61]. While alginate microgels typically exhibit moduli ranging from 0.2 to 20 kPa, depending on the cross-linking agent, the resulting modulus of ∼150 Pa was an intentional design choice, keeping the beads ultra-soft and within the range of tissue-mimetic matrices so as to permit cell spreading, spheroid compaction, and later extracellular-matrix remodeling, while remaining mechanically stable enough to withstand retrieval and handling. The observation that neither In-Bead spheroid formation nor nanoparticle encapsulation measurably altered this modulus is consistent with the low volume fraction of the inclusions relative to the alginate matrix, in which the bulk mechanical response is governed by the alginate network rather than by the embedded spheroids or nanoparticles.

For the first time, we demonstrated that incorporating Nx5.NP into Sph.Beads enhanced cell viability. While a slight viability decrease was observed at Day 1 for both Sph.Beads/Nx5.NP and Sph.Beads/Blk.NP, likely due to high NP loading approaching cellular tolerance thresholds, viability significantly improved by Day 4 in the Nx5.NP group. This supports the protective role of NecroX-5, though its effect might be more pronounced at concentrations closer to 10 µM. However, simply increasing NP content is not viable, as it could compromise early viability and risk altering alginate viscosity or NP stability. Since improving encapsulation efficiency to boost Nx5 payload per NP proved challenging, future work could explore alternative nanocarriers or hydrophilic therapeutics (e.g. RNA-based drugs), where diffusion losses may be less critical. This system thus holds promise for broader applications beyond lipophilic compounds.

## 5 CONCLUSION

By merging biomaterial-based cell engineering with targeted nanotherapeutics, our approach creates a supportive microenvironment that improves SCAP viability over time. This protocol differs from both our previously reported magnetic-bioprinting approach and from other alginate cell-encapsulation methods in three respects. First, flow-focusing droplet generation is continuous and inherently scalable, producing large numbers of beads in a single run without the spatial-resolution and raster-speed constraints of bioprinting. Second, the single-emulsion (water-in-oil) configuration co-encapsulates the NecroX-5-loaded nanoparticles homogeneously throughout the alginate matrix, whereas double-emulsion (water-in-oil-in-water) systems, although effective for forming spheroids in a controlled microenvironment [54], require additional interfacial stabilization and are more sensitive to droplet coalescence during the prolonged 18-h incubation. Third, the short, cell-permissive cross-linking used here preserves the ultra-soft, tissue-mimetic character of the beads while retaining sufficient mechanical integrity for retrieval, a balance that is difficult to achieve with the stiffer, longer cross-linked alginate beads commonly reported for MSC encapsulation [52, 53]. Together, these features make the protocol suitable for the high-throughput generation of functionalized, nanomedicine-loaded MSC spheroids from a standard microfluidic setup. This innovative strategy holds significant promise for improving MSC-based therapies in regenerative medicine.

## Acknowledgements

Floriane Debuisson is a PhD fellow of the Belgian *Fonds National de la Recherche Scientifique* (F.R.S.-FNRS). This work was supported by a F.R.S.-FNRS *Projet de Recherche* (MODULSCI) granted to Anne des Rieux. Ana Santos-Coquillat is grateful for financial support from F.R.S.-FNRS as *Chargée de recherches.* The authors acknowledge Pr. Anibal Diogenes (Dept. Endodontics/UT Health San Antonio), Pr. Thomas Michiels and Stephane Messe (VIRO/De Duve Institute/UCLouvain).

## 6 SUPPLEMENTARY MATERIALS

**Supplementary Video 1:**
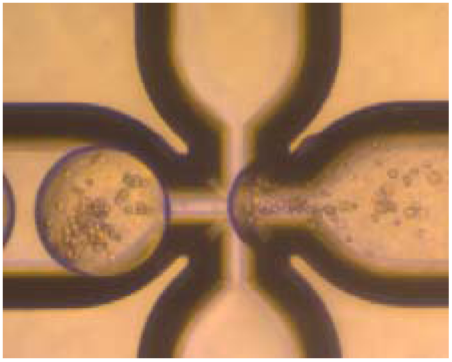
Microfluidic droplet generation. High-speed camera video of the production of droplets using the Dolomite™ microfluidics system with a 190 µm fluorophilic chip. SCAP introduced at 5.10^6^ cells/mL in alginate solution (0.75% -CaEDTA 37.5mM) with Blk.NP with a flow rate of 10 µL/min. Organic phase HFE 7500 with 1% Fluosurf at 20 µL/min.

**Supplementary Video 2:**
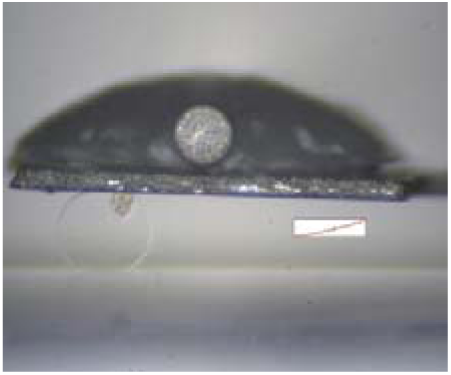
Microsquisher – Sph.Beads/Blk.NP. Images acquired using the Microsquisher^®^ (CellScale). The force required to achieve a 20% displacement was recorded. Bead containing a SCAP spheroid and Blk.NP presented in this video.

